# *PSP2*, a gene encoding RGG-motif protein, is a novel suppressor of clathrin heavy chain deficiency

**DOI:** 10.1101/2020.03.11.986810

**Authors:** Debadrita Roy, Mani Garg, Purusharth I Rajyaguru

## Abstract

Clathrin, made up of the heavy- and light-chains, constitutes one of the most abundant protein in vesicles involved in intracellular protein trafficking and endocytosis. *YPR129W*, which encodes RGG-motif containing translation repressor was identified as a part of multi-gene construct (SCD6) that suppressed clathrin deficiency. However, the contribution of *YPR129W* alone in suppressing clathrin deficiency has not been documented. In this study we identify *YPR129W* as a necessary and sufficient gene in a multigene construct SCD6 that suppresses clathrin deficiency. Importantly, we identify cytoplasmic RGG-motif protein encoding gene *PSP2* as a novel suppressor of clathrin deficiency. Three other RGG-motif protein encoding genes *SBP1, DED1* and *GBP2* do not suppress clathrin deficiency. *DHH1*, a DEAD-box RNA helicase with translation repression activity also fails to rescue clathrin deficiency. α-factor secretion assay suggests that suppression of clathrin deficiency by *SCD6* and *PSP2* is not mediated by the rescue of the trans-Golgi network (TGN) protein sorting defect observed in the absence of *CHC1*. Detailed domain analysis of the two suppressors reveals that the RGG-motif of both Scd6 and Psp2 is important for suppressing clathrin deficiency. Additionally, the Lsm domain deletion as well as the arginine to alanine mutation in the arginine methylation defective (AMD) mutant render Scd6 defective in suppressing clathrin deficiency. Overall based on our results using *SCD6* and *PSP2* proteins, we identify a novel role of RGG-motif in suppressing clathrin deficiency. Since both the suppressors are RNA-binding granule-resident proteins, this study opens an exciting avenue for exploring the connection between clathrin function and cytoplasmic RNA metabolism.

## Introduction

Clathrin is a protein of vital importance in cells. Clathrin coated vesicles (CCVs) play a key role in endocytosis and various intracellular trafficking. The triskelion, which is made up of the clathrin light chain and heavy chain monomers, forms a typical polygonal surface lattice on CCVs. Clathrin mediated endocytosis, which is highly conserved from yeast to humans, is the major pathway for the internalization of external and membrane molecules into the cell. There are clathrin-independent pathways, however without clathrin-mediated endocytosis, uptake is greatly reduced ^1–4^. CCVs are also involved in the intracellular trafficking pathway. Deficiency of clathrin in unicellular organisms like yeast, amoebae and protozoa are associated with defects in intracellular trafficking and receptor-mediated endocytosis ^5,6^.

In certain yeast strains, the deletion of the clathrin heavy chain-1 gene (*CHC1*) renders cells inviable, whereas in others, cells are viable. This is owing to the presence of an independently segregating gene – suppressor of clathrin deficiency 1 (*SCD1).* Presence of the *scd1-i* allele leads to lethality in *chc1* deficient cells, whereas the *scd1-v* allele allows growth, albeit poorly. Along with slow growth, chc1Δ scd1-v cells have abnormal morphology, genetic instability, reduced endocytosis and mis-localization of endoproteases from the *trans*-Golgi network to the cell membrane ^7–12^. *SCD1* was recently identified to encode *PAL2*, a protein involved in localizing to cortical patches with other endocytic factors, thereby facilitating endocytosis. It was found that scd1-i allele has a premature stop codon which results in a truncated version of wild type *PAL2*, encoded by *scd1-v* allele ^13^. YEpSCD6 was one of the six multi-gene multicopy suppressors identified in a genetic screen ^14^. This suppressor contains a total of 8 genes along with *YPR129W*. A personal communication on *Saccharomyces* genome database indicated *YPR129W* as a multicopy suppressor of clathrin deficiency (Gelperin et al. 1995, personal communication to SGD). *YPR129W* encodes an RGG-motif translation repressor protein that targets translation initiation factor eIF4G1 ^15^. We wanted to confirm its role in affecting clathrin function and further address the possible role of other RGG-motif containing RNA-binding proteins in affecting clathrin function. We hypothesized that other RGG-motif proteins could also affect clathrin function.

RGG-motif proteins contain single or multiple repeats (Table 1) of RGG/RGX (‘X’ being any residue). This motif is known to be involved in protein-nucleic acids and protein-protein interactions ^16^. The RGG-motif containing proteins function in processes like transcription, cell cycle, apoptosis and synaptic plasticity. The RGG-motif constitutes the second most common class of domains/motifs that contribute to RNA-binding ^17^. Recently, a subset of yeast RGG-motif proteins (such as Scd6, Sbp1, Npl3 and Ded1) have been implicated in translation control ^15,18^ as these proteins bind conserved translation initiation factor eIF4G through their RGG-motif to repress translation. Scd6 and Sbp1 are translation repressors and decapping activator proteins. Consistent with their role, both Scd6 and Sbp1 localize to RNA granules that are sites of translation repression and mRNA decay ^15,19,20^. *PSP2* encodes a cytosolic RGG-motif (Table 1) containing protein originally identified as a suppressor of intron-splicing defects of a mutation in *MRS2* and subsequently as a suppressor of conditional mutation in DNA POL I in yeast ^21,22^. Interestingly, Psp2 localizes to processing bodies during glucose deprivation ^19^. *DED1* encodes a DEAD-box RNA helicase and a RNA granule resident protein that can both promote and inhibit translation initiation in yeast ^23,24^. *GBP2* encodes a nucleo-cytoplasmic shuttling protein involved in telomere length modification ^25^, mRNA export ^26–28^ and mRNA quality control ^29^. Despite being a nuclear protein, Gbp2 associates with polysomes and is a component of stress granules ^30,31^ the significance of which remains to be understood. *DHH1* encodes a DEAD-box RNA helicase that acts as a translation repressor and decapping activator ^32^. It localizes to RNA granules and regulates P-body assembly in ATP-dependent manner ^33^. In this work, we report that *YPR129W* is a bonafide suppressor of clathrin deficiency. We identify *PSP2*, a gene encoding RNA-binding and RGG-motif protein as a novel suppressor of clathrin deficiency. The RGG-motif of both Scd6 and Psp2 is important for suppression of clathrin deficiency. These results establish an unprecedented link between clathrin heavy chain function and granule-resident RGG-motif proteins. Our observations pave the way for exploring the mechanistic basis underlying this link in the future.

**Table 1:** Number of RGG/RGX in different proteins tested in this study

| Protein (# of amino acids) | # of RGG-/RGX- | Location of RGG-/RGX- motifs |
| --- | --- | --- |
| Scd6 (349) | 1/7 | C-terminal |
| Psp2 (593) | 4/10 | C-terminal |
| Sbp1 (294) | 8/5 | Central |
| Ded1 (604) | 4/4 | Dispersed |
| Gbp2 (427) | 4/6 | Predominantly towards N-terminal |

## RESULTS

### *YPR129W* is necessary and sufficient to suppress clathrin deficiency

YEpSCD6, a high copy number multi-gene plasmid was reported to be capable of rescuing inviability of clathrin heavy chain-deficient yeast strain expressing endogenous *CHC1* under a galactose-inducible promoter ^14^. This multi-gene construct contained 7 other genes along with *YPR129W.* Based on this study and the personal communication (Gelperin et al. 1995, personal communication to SGD), we were intrigued by the possible role of an RNA-binding translation repressor protein in suppressing clathrin heavy chain requirement and therefore wanted to indeed confirm the role of *YPR129W* in clathrin function. To test in our hands, the necessity and sufficiency of *YPR129W*, we created YEpΔ*SCD6* (lacking *YPR129W*) and YEponly*SCD6* (YEpo*SCD6*; expressing only *YPR129W*) by site directed mutagenesis. These plasmids (Table 3) were transformed into *GAL1:CHC1* strain and the transformants were assayed for growth on glucose and galactose. We observed that YEpΔ*SCD6* transformant grew as poorly as the empty vector control on glucose plate (Figure 1A) indicating that in the absence of *SCD6* none of the other 7 genes could suppress the clathrin deficiency growth defect. Thus, *SCD6* was necessary in the multi-gene construct to suppress clathrin deficiency mediated growth defect. YEpo*SCD6* transformed cells grew in manner comparable to YEp*SCD6* transformed cells suggesting that *SCD6* was sufficient to suppress the clathrin deficiency mediated growth defect (Figure 1A). In order to quantify the extent of rescue, a plating assay was carried out on YEP media, wherein after depleting Chc1, cells were plated on galactose and glucose followed by counting the colonies. Consistent with the growth assay, cells expressing YEp*SCD6* and YEpo*SCD6* but not YEp*Δscd6* could suppress clathrin deficiency. (Figure 1B). To further confirm its role, we cloned *YPR129W* in a 2u plasmid pGP564 with a C-terminal His-tag and checked its ability to suppress clathrin deficiency. We observed that this plasmid encoded *SCD6* suppressed the clathrin deficiency-mediated growth defect in a manner comparable to YEp*SCD6* (Figure 1C). Based on these results we confirmed and established that *YPR129W* is a suppressor of clathrin deficiency.

**Table 3:**
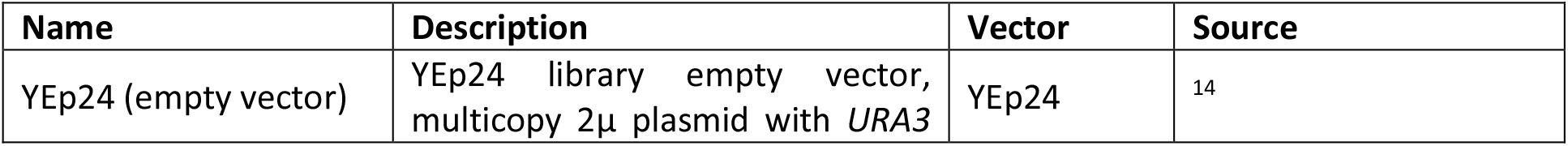
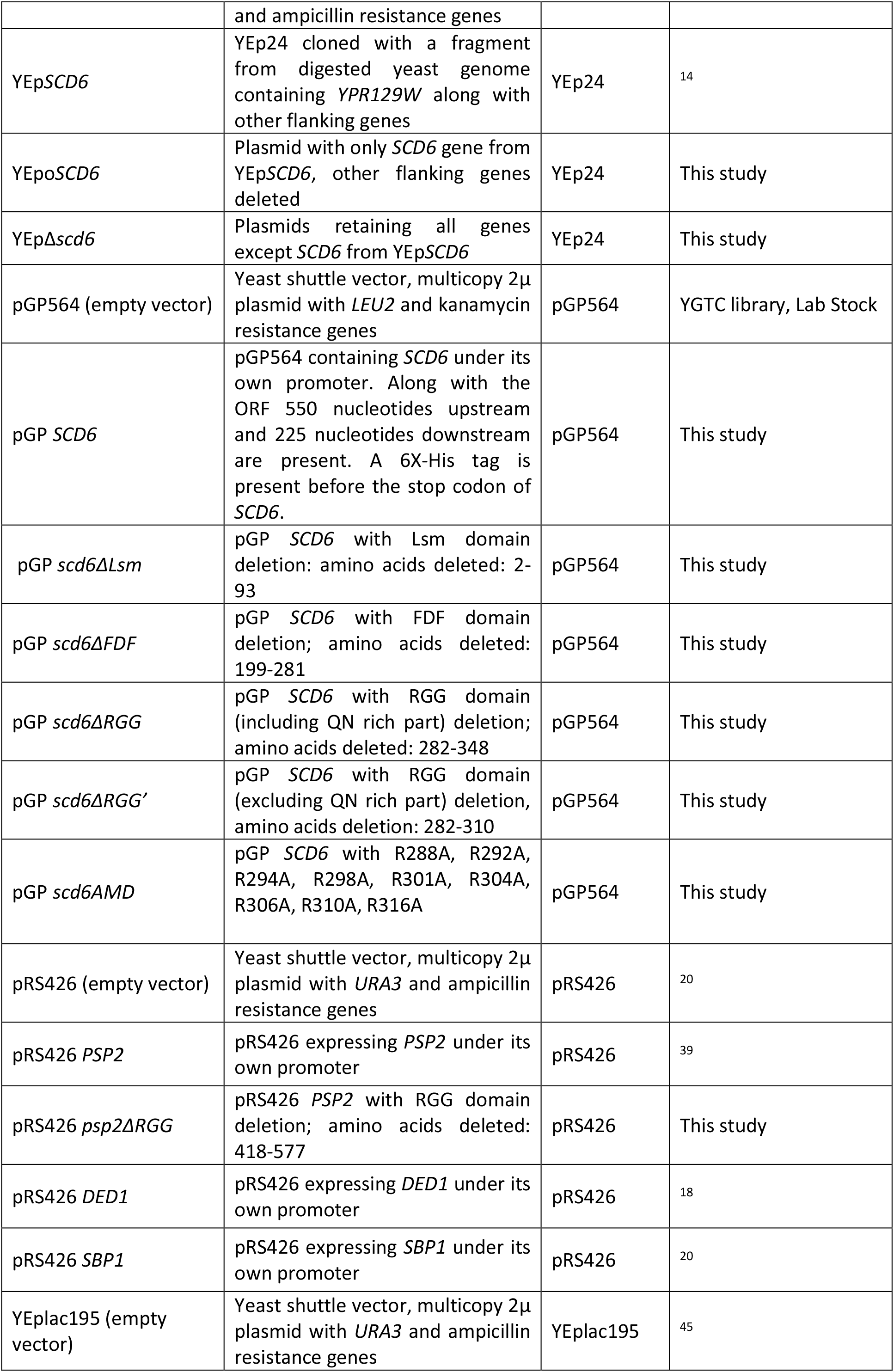
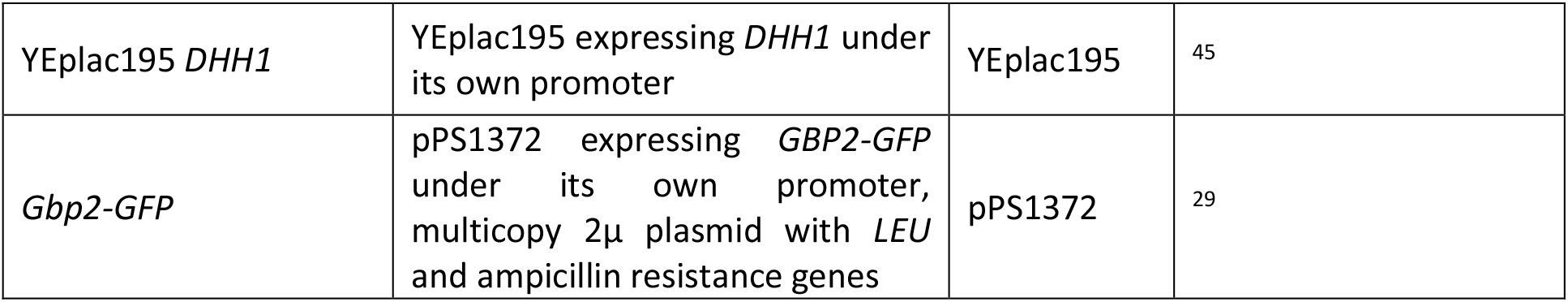
List of plasmids used in this study

**Figure 1:**
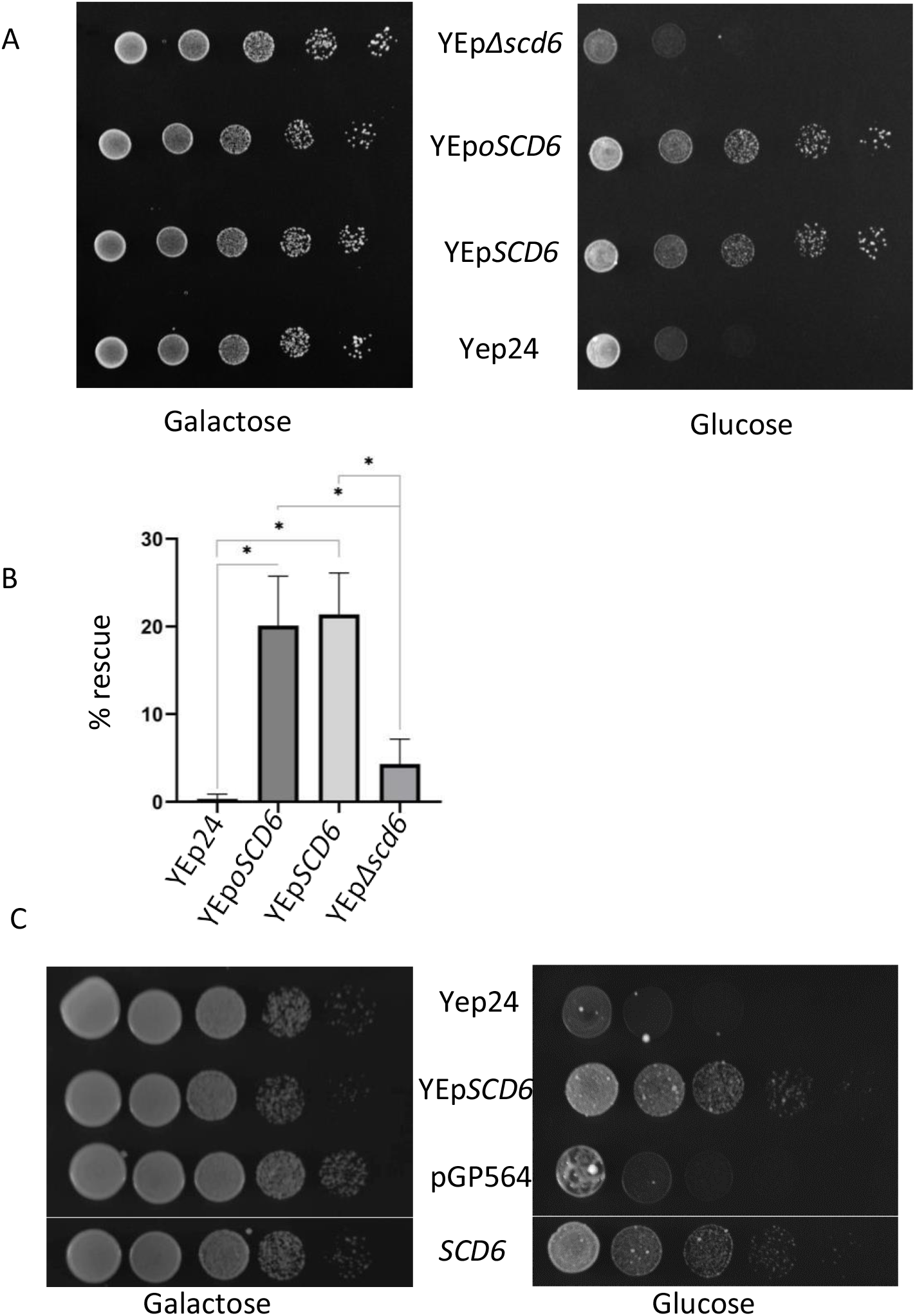
*YPR129W* is necessary and sufficient to rescue clathrin deficiency. (A) Growth assay of *GAL1:CHC1* cells (endogenous *CHC1* is under galactose-inducible promoter) transformed with YEp24 (empty vector), YEp*SCD6*, YEp*Δscd6* (lacking *YPR129W*) and YEp*oSCD6* (YEponly*SCD6* expressing only *YPR129W*). 10 OD_600_ cells were serially diluted and spotted on glucose or galactose containing selective media plates. These plates were incubated at 30°C for 2 days (galactose media) or 4-5 days (glucose media) before imaging. (B) Quantification of the percentage rescue of *GAL1:CHC1* transformants upon clathrin deficiency (* indicates p<0.05). Percentage rescue calculated as (number of colonies on glucose/number of colonies on galactose) *100. (C) Growth assay of *GAL1:CHC1* transformants. *SCD6* panel refers to the transformant expressing *SCD6* from pGP564 vector with C-terminal His-tag. *SCD6* expressed from pGP564 vector was spotted on the same plate as the rest of the three spottings. Image was spliced to remove other spottings not relevant to this figure. (Note: Because of the difference between the selection marker between Yep and pGP564, this assay was done on YEP plates). Plates were incubated at 30 °C for 2 days (galactose media) or 4-5 days (glucose plate) before imaging.

### *PSP2* but no other RGG-motif protein encoding genes can suppress of clathrin deficiency defect

*SCD6* encodes RGG-motif containing protein which acts as a translation repressor. We wanted to check whether other genes encoding RGG-motif proteins and/or translation repressors can suppress clathrin deficiency. We therefore tested *DED1*, *SBP1*, *DHH1*, *GBP2* and *PSP2*. Sbp1, Ded1 and Dhh1 can act as translation repressor proteins. Sbp1 and Ded1 (but not Dhh1) contains RGG-/RG-motifs which vary in number of repeats and location in the protein sequence (Table 1). Gbp2 is a shuttling RGG-motif protein which predominantly localizes to the nucleus. Psp2 is an RGG-motif containing protein, which localizes to RNA granules and has recently been reported to modulate translation of specific mRNAs involved in autophagy (Yin et al., 2019). Interestingly *DED1*, *SBP1*, *DHH1* and *GBP2* did not suppress the clathrin deficiency mediated growth defect (Figure 2A). Plating assay further confirmed the growth assay results (Figure 2B). Strikingly, *PSP2* suppressed clathrin deficiency mediated growth defect (Figure 3A). Plating analysis followed by colony count confirmed the results obtained using growth assay (Figure 3B). The percentage rescue by *PSP2* improved when the plating assay was performed on selective synthetic media plates likely due to increased retention of the plasmid (Supp. Figure 1). Based on these results we conclude that *PSP2* is a novel suppressor of clathrin deficiency. These results also indicate that the genes encoding RGG-motif containing or translation repressor proteins in general do not suppress clathrin deficiency highlighting the specificity of suppression by *SCD6* and *PSP2*.

**Figure 2:**
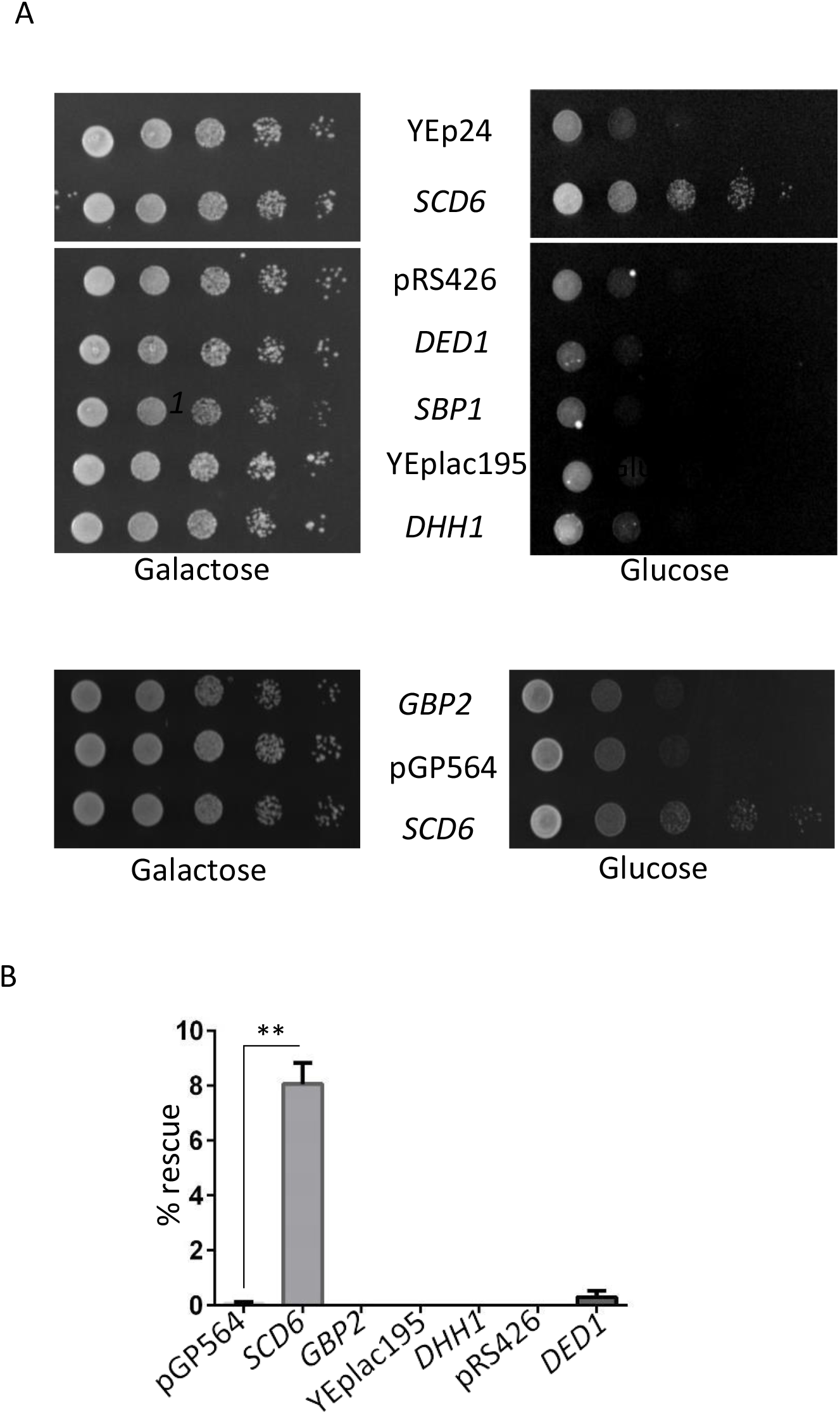
*DED1*, *SBP1*, *DHH1* and *GBP2* fail to rescue clathrin deficiency growth defect. A) Growth assay of *GAL1:CHC1* cells transformed with different plasmids as labeled. 10 OD_600_ cells were serially diluted and spotted on glucose or galactose containing selective media plates. These plates were incubated at 30°C for 2 days (galactose media) or 4-5 days (glucose media) before imaging. YEp24 and *SCD6* transformed cells were spotted on the same plate as the rest of the spottings. B) Quantification of plating assay to measure percentage rescue by *GBP2*, *DED1* and *DHH1* (** indicates p<0.01). Percentage rescue was calculated as the number of colonies on glucose plate/number of colonies on galactose plate*100.

**Figure 3:**
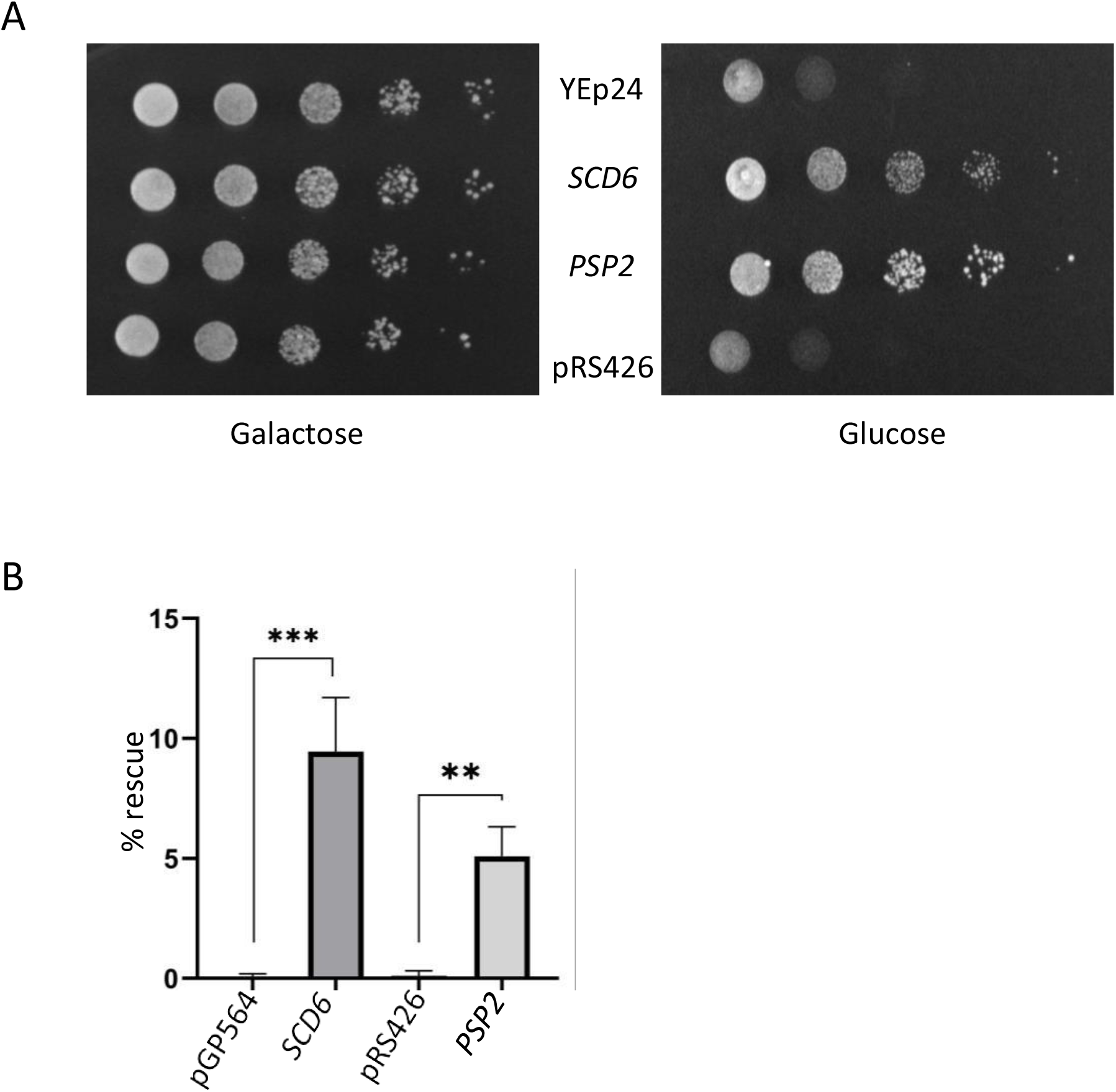
*PSP2* is a novel suppressor of clathrin deficiency. (A) *GAL1:CHC1* cells were transformed with pRS426 (empty vector) or pRS426-*PSP2.* 10 OD_600_ cells were serially diluted and spotted on glucose or galactose containing selective media plates. These plates were incubated at 30°C for 2 days (galactose media) or 4-5 days (glucose media) before imaging. (B) Quantification of the percentage rescue by *PSP2* upon clathrin deficiency using plating assay (** and *** indicates p<0.01 and p<0.001 respectively). Percentage rescue was calculated as the number of colonies on glucose plate/number of colonies on galactose plate*100.

### *SCD6* and *PSP2* do not rescue the trans-Golgi network (TGN) sorting function of the *CHC1*

We next tested the ability of *SCD6* and *PSP2* to rescue the TGN sorting defects of *CHC1* depleted cells. MATα cells are defective in the secretion of mature form of the mating pheromone α-factor, in the absence of clathrin. This is due to the mis-localization of α-factor processing enzymes from the trans-Golgi network to the plasma membrane ^9^. As a result, MATα cells with defective clathrin function do not inhibit the growth of MATa cells otherwise evident by a zone of growth inhibition (halo) in the halo assay. Since *SCD6* and *PSP2* rescue the growth defect phenotypes upon depletion of Chc1, we decided to test their contribution to α-factor secretion. *GAL1:CHC1* MATα cells overexpressing plasmid encoding *CHC1*, *SCD6*, *SBP1*, *PSP2*, *GBP2* or *DED1* were depleted of Chc1 by growing on glucose and spotted on a lawn of MATa cells expressing the *sst1-2* allele (BJ3556 strain in Table 2). *GAL1:CHC1* MATα cells overexpressing *CHC1* from a plasmid form a clear zone of growth inhibition (halo) however such a zone of inhibition is absent for cells overexpressing any of the tested genes including *SCD6* and *PSP2* (Figure 4). We interpret this result to suggest that the suppression of clathrin deficiency growth defect by *SCD6* and *PSP2* does not involve complementation of the TGN function.

**Table 2:**
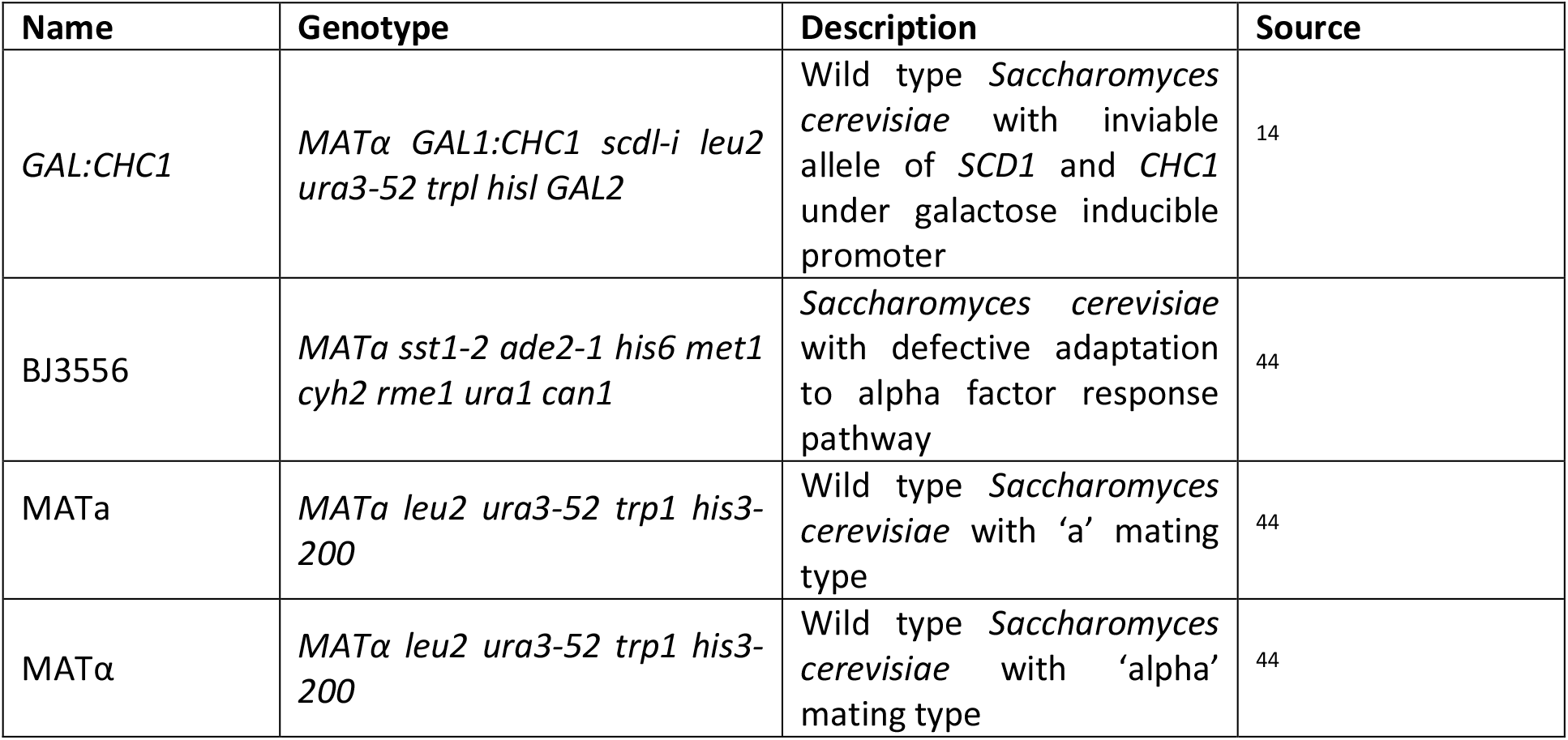
List of strains used in this study

**Figure 4:**
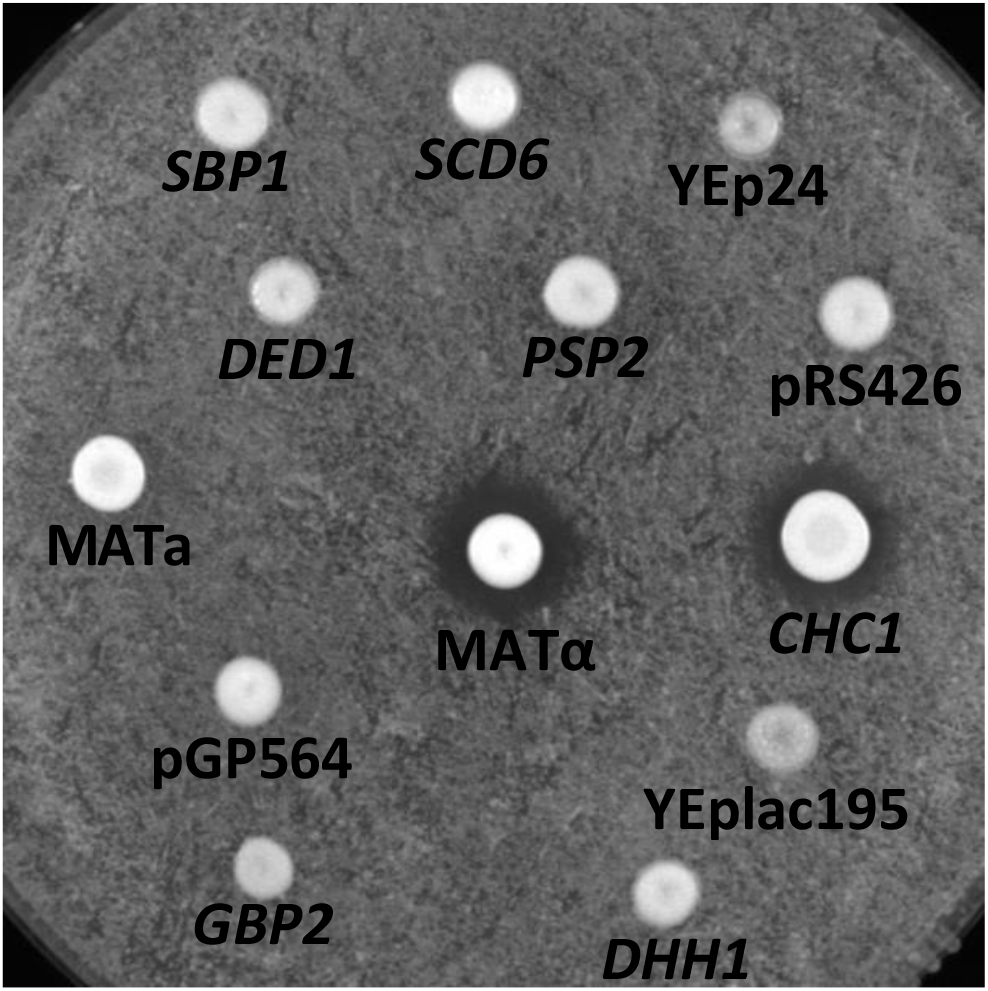
*SCD6* and *PSP2* fail to rescue the trans-Golgi network (TGN) sorting function of clathrin heavy chain. MATα strain expressing endogenous *CHC1* under galactose-inducible promoter was transformed with plasmids expressing *SCD6, SBP1, DED1, PSP2, GBP2, DHH1* or *CHC1*. Respective empty vectors were used as control. Transformants were spotted on a lawn of MATa cells carrying the *sst1-2* allele and incubated on YEP glucose plate until (2-3 days) the appearance of a zone of clearance (halo). YEp24 is the empty vector control for *SCD6* and *CHC1*. pRS426 is the empty vector control for *DED1, SBP1* and *PSP2*. pGP564 is the empty vector control for *GBP2* and YEplac195 is the empty vector control for *DHH1*. Wild type MATa and MATα strains act as negative and positive controls,respectively.

### RGG-motif plays a vital role in suppressing clathrin-deficiency

Both *SCD6* and *PSP2* contain RGG-motif rich C-terminal region. *SCD6* also contains an Lsm domain at its N-terminus and a central FDF domain. *PSP2* does not contain any other canonical domain/motif other than the RGG-motif. We investigated which domain of these proteins was responsible for the observed suppression of growth defect. To this end, we generated domain deletion constructs and assayed them for suppression of clathrin deficiency mediated growth defect. An RGG-deletion mutant of *PSP2* was created and tested for its ability to suppress clathrin deficiency. We observed that the RGG-motif deletion mutant of *PSP2* was defective in suppressing clathrin deficiency (Figure 5A). Plating assays followed by colony counting further confirmed that the RGG-motif deletion mutant of *PSP2* was defective in rescuing growth on the glucose media (Figure 5B and Supp. Figure 1). We conclude that the RGG-motif of *PSP2* plays an important role in suppressing clathrin deficiency.

**Figure 5:**
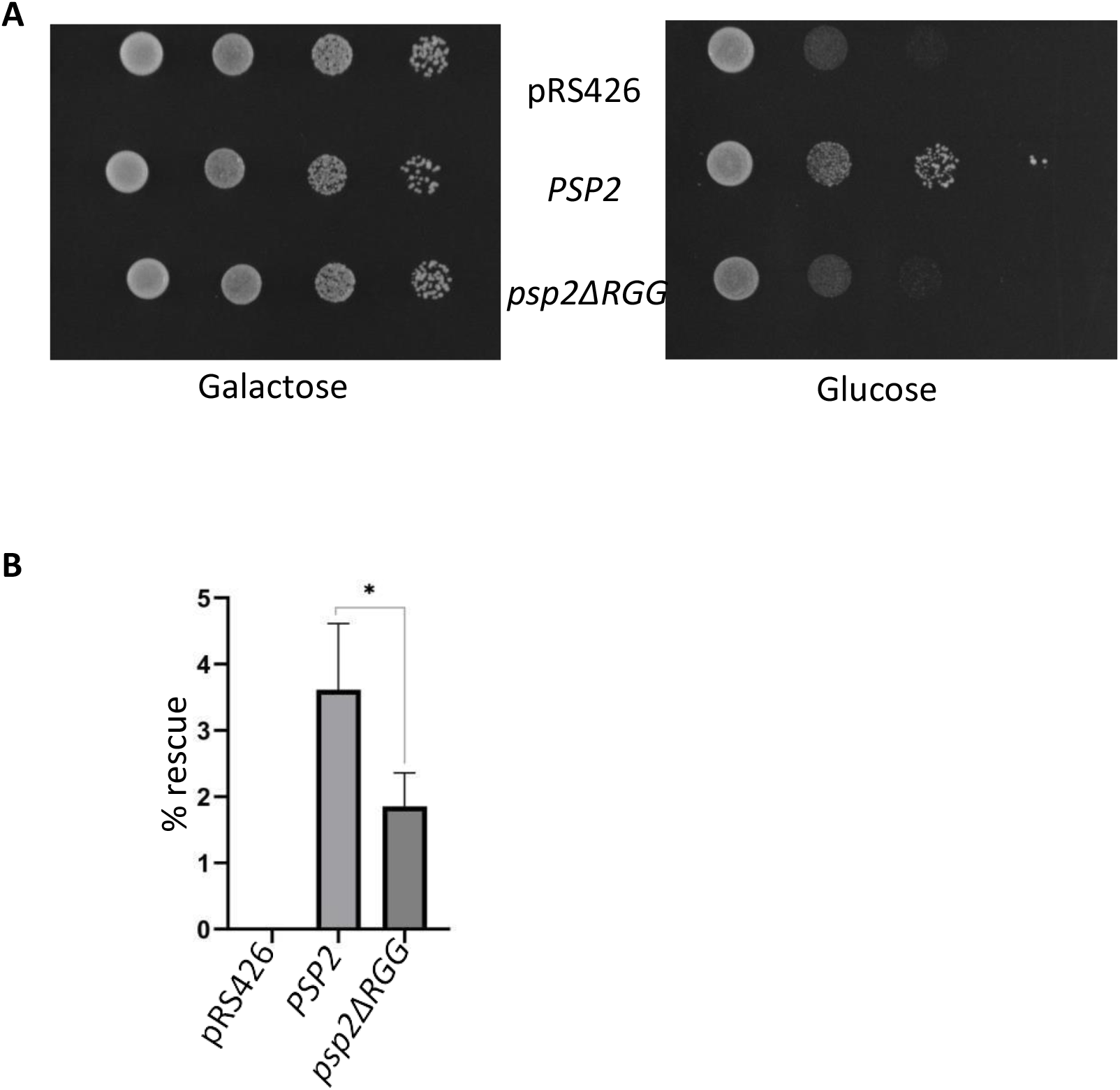
Deletion of RGG-motif compromises the ability of Psp2 to suppress clathrin deficiency growth defect. (A) Growth assay of *GAL1:CHC1* cells transformed with wild type and RGG-motif deletion mutants of Psp2. 10 OD_600_ cells were serially diluted and spotted on glucose or galactose containing selective media plates. These plates were incubated at 30°C for 2 days (galactose media) or 4-5 days (glucose media) before imaging. (B) Quantification of the percentage rescue of *GAL1:CHC1* transformants with wild type or RGG-motif deletion mutant of PSP2 upon clathrin deficiency (* indicates p<0.05). Percentage rescue was calculated as the number of colonies on glucose plate/number of colonies on galactose plate*100.

The RGG-motif deletion construct of *SCD6* was also highly defective in suppressing clathrin deficiency (Figure 6A). The Lsm domain deletion construct was also defective in suppressing clathrin deficiency albeit to slightly lesser extent than the RGG-motif deletion construct. Deletion of the FDF motif weakly affected the ability of the mutant to suppress clathrin deficiency. The RGG-motif (282-348 residues) of *SCD6* as annotated in literature ^34^ comprises of RGG-rich sequences (282-310 residues) and QN-rich sequences (311-348 residues). Deletion of just the RGG-rich sequences (282-310) (*scd6RGG’*) resulted in a defect that was comparable to the deletion of 282-348 residues indicating that the RGG-rich sequence was indeed important for the rescue of clathrin deficiency. Based on these results we conclude that all the three domains affect the ability of *SCD6* to suppress clathrin deficiency. RGG-motif is the least dispensable motif and FDF-domain is the most dispensable domain for suppressing clathrin deficiency. Plating experiments followed by colony counting confirmed the growth assay results (Figure 6B). Interestingly the FDF domain deletion mutant was also defective in suppressing clathrin deficiency in the plating assay which was not evident as much in the growth assay (Figure 5A). The differential behavior of this mutant in two different assays is interesting however the basis for this is unclear. Some of the arginine residues in the RGG-motif of Scd6 are methylated and this modification promotes the repression activity ^35^. We tested the role of arginine residues of the RGG-motif in suppressing clathrin deficiency. We observed that the arginine methylation defective mutant (AMD; with 9 R to A substitutions) failed to suppress the clathrin deficiency both in growth and plating assay (Figure 6C & D). It is possible that arginine methylation of *SCD6* RGG-motif could play a role in suppression of clathrin deficiency. Endogenous Scd6 is expressed in low copy number [1280 molecules/cell ^36^] and we believe that its expression is highly regulated. Detection of Scd6 and its mutants expressed from pGP564 (Figure 6) using an anti-His antibody has been technically challenging. We however know that expression of both RGG-deletion and AMD mutant of Scd6 is not compromised when expressed under galactose-inducible promoter from a 2u plasmid (BG1805 vector) and detected using PAP reagent that detects the C-terminal ZZ-tag ^35^. We also know that expression of Lsm domain deletion mutant is not compromised when expressed in the BG1805 vector (Parbin et al., under review). Overall these results indicate that the Scd6 RGG-motif and arginines in RGG-motif are important for suppression of clathrin deficiency. Interestingly the Lsm and the FDF (to a certain extent) domains also contribute to suppression of clathrin deficiency by Scd6.

**Figure 6:**
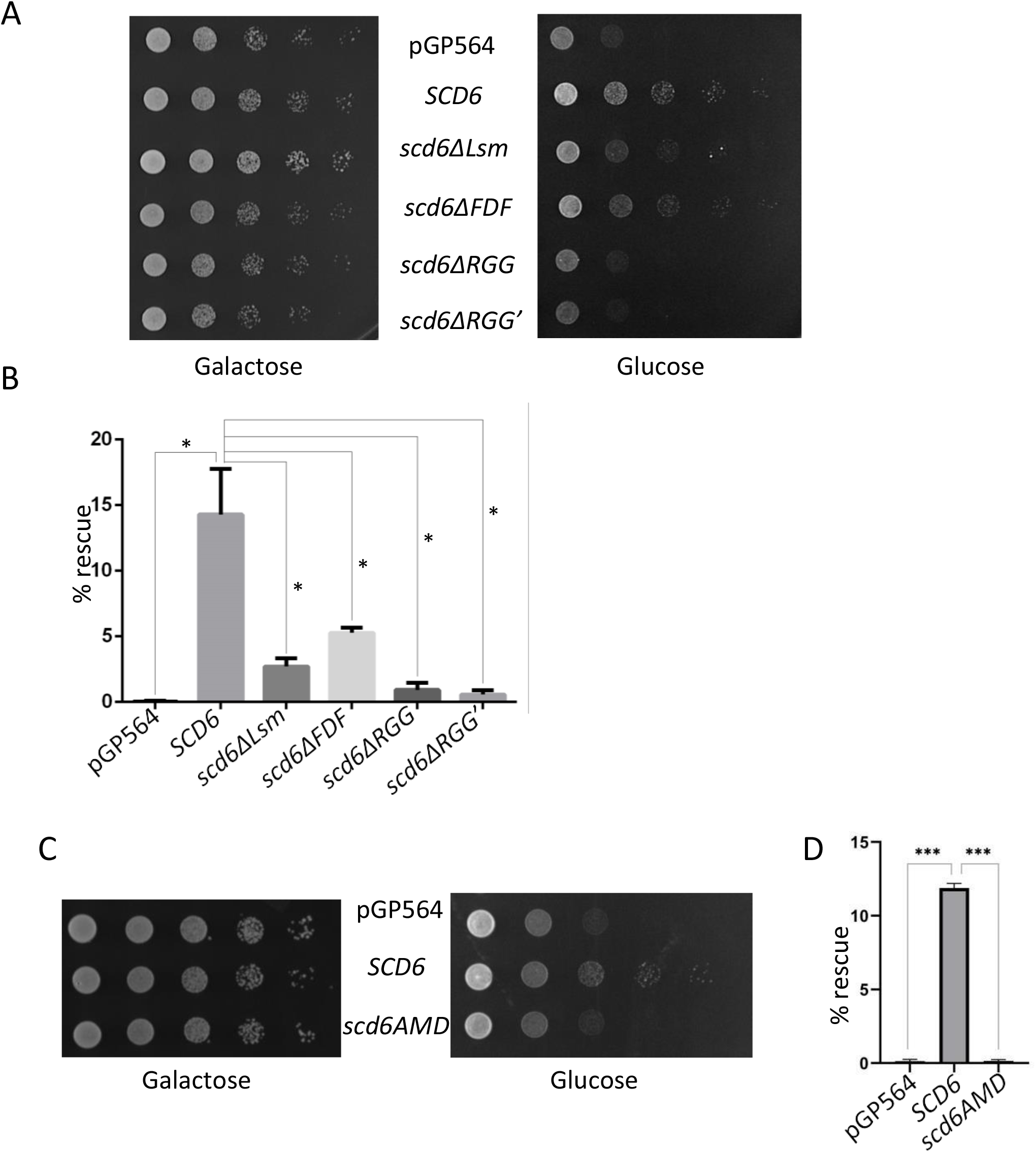
RGG-motif and Lsm domain are important for suppress clathrin deficiency growth defect by Scd6. (A) Growth assay of *GAL1:CHC1* cells transformed with plasmids expressing wild type, Lsm, FDF, RGG or RGG’ domain/motif deletion mutants of Scd6. 10 OD_600_ cells were serially diluted and spotted on glucose or galactose containing selective media plates. These plates were incubated at 30°C for 2 days (galactose media) or 4-5 days (glucose media) before imaging. (B) Quantification of the percentage rescue of *GAL1:CHC1* by *SCD6* mutants upon clathrin depletion. (C) Growth assay to test the ability of arginine methylation defective (AMD) mutant of *SCD6* to suppressing clathrin deficiency. (D) Quantification of the percentage rescue of *GAL1:CHC1* transformants by the AMD mutant of *SCD6* upon clathrin deficiency (* and ** indicates p<0.05 and p<0.01 respectively). Percentage rescue was calculated as the number of colonies on glucose plate/number of colonies on galactose plate*100.

## Discussion

We provide evidence suggesting that two genes encoding RGG-motif proteins function as suppressors of clathrin deficiency. To the best of our knowledge this is the first peer-reviewed report implicating RGG-motif proteins in clathrin function. Our conclusion is based on the following observations, a) *SCD6* acts as a suppressor of clathrin deficiency (Figure 1) b) *PSP2* is a novel suppressor of clathrin deficiency (Figure 3) c) RGG-motifs of Psp2 and Scd6 are required for suppressing clathrin deficiency (Figure 5 and 6) d) Arginine residues of the Scd6 RGG-motif play an important role in suppressing clathrin deficiency (Figure 6C and D).

*SCD6* was identified as one of the genes in a multi-gene construct that suppressed clathrin heavy chain deficiency, however the specific contribution of *SCD6* remained to be confirmed. Our results prove that in the reported multi-gene construct, *SCD6* is both necessary and sufficient to suppress clathrin deficiency mediated growth defect. This along with the observation that *SCD6* cloned in a different vector also suppresses clathrin deficiency indicates that it is a genuine suppressor of clathrin deficiency. Reported roles of *SCD6* orthologs point towards a role of this family of proteins in endocytosis and/or trafficking. *CAR-1* (worm ortholog) plays a role in maintaining ER organization as upon *CAR-1* knockdown, ER is disorganized into large patches and thick strands ^37^. Trailer hitch (Drosophila ortholog) localizes to ER-exit sites, which persist as large foci upon Tral knockdown ^38^. Whether *SCD6* functions along similar lines as its worm and fly ortholog is unclear.

Suppression of clathrin deficiency by *PSP2* is a striking result. Psp2 is known to localize to RNA granules ^19^ and upon overexpression, can rescue P-body formation in *edc3Δlsm4Δc* strain ^39^. Scd6 binds *TIF4631* (eIF4G1) to repress translation and interestingly Psp2 is reported to physically interact with its paralog *TIF4632* ^40^. A recent report has indeed implicated Psp2-TIF4632 interaction in modulating translation of autophagy genes (Yin et al., 2019). Whether Psp2 can act as a translation repressor of specific mRNAs has not been explored. Requirement of the Psp2 RGG-motif in suppressing clathrin deficiency (Figure 5B) points toward a mechanism likely to be similar to that of Scd6 which needs to be tested further.

Lack of suppression by other RGG-motif proteins and translation repressors (Sbp1, Ded1, Dhh1 and Gbp2) highlights the specificity of Scd6 and Psp2 in clathrin function. It is unlikely that the contribution of Scd6 and Psp2 in clathrin function is dependent on the number of RGG-/RGX-repeats as Sbp1 contains more repeats than Scd6 (Table 1) but fails to suppress. Interestingly both Scd6 and Psp2 harbor RGG-motifs at its C-terminal but none of the other tested proteins including Sbp1 have RGG-motifs at their C-terminus.

Clathrin deficiency suppression mechanisms might likely fall into one of the following two categories: a) Suppressors that directly contribute to clathrin function in vesicular transport/endocytosis and b) Suppressors that indirectly increase tolerance to clathrin deficiency. *SCD4* ^41^ and *SCD5* ^11^ belong to the former category that contribute directly to vesicular transport. *SCD2* on the other hand belongs to latter category of suppressors. It was identified as *UBI4* ^14^ which likely suppresses clathrin deficiency defect by accelerating clearance of mis localized proteins We hypothesize that *SCD6* and *PSP2* might act in an indirect manner by altering translation to regulate the load of newly synthesized proteins on the trafficking machinery. This hypothesis is based on the following observations: 1) RGG-motif, which is required for translation repression activity of *SCD6* 15,35,42 is also required for suppressing clathrin deficiency defect by *SCD6* and *PSP2* 2) *SCD6* AMD mutant is defective in repressing translation as well as in suppressing clathrin deficiency 3) Lsm domain is required for the translation repression activity of Scd6 (Parbin et al., *communicated*) and also for suppressing clathrin deficiency. Several key experiments will be required in future to test this hypothesis. It is also possible that translation control of specific mRNAs involved in endocytosis and/or trafficking by Scd6 and perhaps Psp2 contributes to the suppression of clathrin deficiency. Identification of specific mRNA targets of Scd6 and Psp2 will provide important insights in this regard. Since Scd6 localizes to RNA granules in RGG-motif dependent manner and RNA granules are sites of translation repression and mRNA decay, we tested the localization of Scd6 upon clathrin depletion. We did not observe any significant difference in localization of Scd6 to foci (data not shown) suggesting that suppression is likely not mediated by changes in localization to foci.

LSM14 is the human ortholog of Scd6 ^34,43^. The role of Scd6 identified in mRNA fate determination in yeast is conserved in humans. It is possible that LSM14 could contribute to clathrin mediated endocytosis and trafficking. This exciting possibility will be tested in future. Overall our results establish a new link between RGG-motif containing RNA-binding proteins and clathrin function.

Analyzing the mechanistic basis of this link could lead to unraveling of the role of mRNA translation control pathway in clathrin mediated endocytosis. It will also be pertinent to test if the players contributing to endocytic/trafficking pathway including Chc1 are involved in modulating cytoplasmic mRNA fate. Our current work thus highlights several interesting research questions and addressing these questions will be an exciting future endeavor.

## Acknowledgements

This work was predominantly supported by Department of Science and Technology grant (EMR/2017/001332) from the government of India. It was also supported by India Alliance DBT-Wellcome trust (IA/I/12/2/500625) and DBT-IISc partnership program (BT/PR27952-INF/22/212/2018). DR was supported by DST-INSPIRE fellowship; MG is supported by University Grants Commission (UGC) fellowship. We are indebted to Sandra Lemmon (University of Miami) for helping us with various strains and reagents. We thank her for providing inputs at different stages of this work. We also sincerely thank Roy Parker (University of Colorado, Boulder) and Kenji Irie (University of Tsukuba) for sharing plasmids and strains.

## Contributions

PIR and DR designed the experiments; DR and MG performed the experiments; PIR, DR, and MG prepared the figures; PIR wrote the manuscript; PIR and MG revised the manuscript.

## Materials and methods

### Strains and Plasmids

Strains and plasmids used in this study are enlisted in Table 2 and Table 3 respectively.

### Growth Assay

Growth assays were performed to assess the ability of plasmids expressing desired genes to rescue clathrin deficiency mediated growth defect of the *GAL:CHC1* strain on glucose media. In *GAL:CHC1* strain, *CHC1* is under a galactose inducible promoter ^14^ which results in a growth defect on glucose media. Transformants grown on galactose containing media (10 OD_600_) were, serially diluted and spotted on the respective minimal media containing glucose and galactose plates. All the plated were incubated at 30°C and imaged after 2 days (galactose media) or 4-5 days (glucose media). Growth assays were performed at least three times with comparable results.

### Plating Assay

After growing cells in YEP-glucose for more than 16 −20h, different *GAL:CHC1* transformants were diluted to 0.001 OD_600_. Three different volumes (50ul, 100ul and 200ul) were plated on YEP-galactose and glucose plates. The galactose containing plates were incubated for 4 days and glucose plates for 7 days before imaging. Colonies were counted and plotted as percentage rescue which was calculated as [(no. of colonies on glucose/no. of colonies on galactose) *100]. Plating assays were performed at least three times with comparable results which were quantitated.

### Halo Assay

The background tester MATa strain (BJ3556) carrying the *sst1-2* allele and the different *GAL:CHC1* transformants were grown on YEP-glucose for 10-12 h and 16-20 h respectively. The tester strain was diluted to 0.1 OD_600_ and spread onto YEP-glucose plate. Subsequently different *GAL:CHC1* transformant cultures were spotted onto the plate which were incubated for 2-3 days before imaging. MATa and MATα strains labelled in the Figure 4 are wild type strain controls.

