## Supplementary Figure 1 for "*PSP2*, a gene encoding RGG-motif protein, is a novel suppressor of clathrin heavy chain deficiency"

% rescue

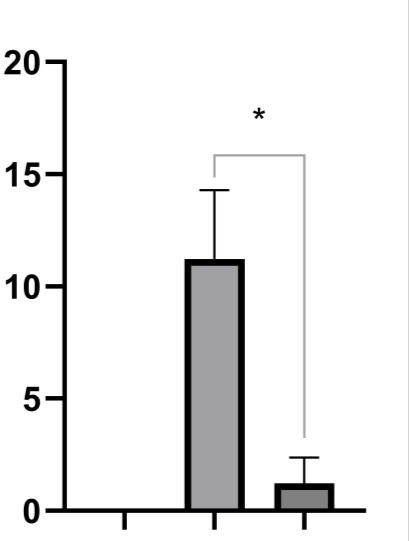

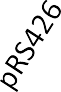

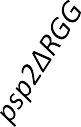

### Supplementary Figure 1: Deletion of the RGG-motif compromises the ability of *PSP2* to suppress clathrin deficiency growth defect. Plating assay performed on selective synthetic media plate followed by quantification of the percentage rescue of different *GAL1:CHC1* transformants (p<0.05). Percentage rescue was calculated as the number of colonies on glucose plate/number of colonies on galactose plate*100.
